# Fine-mapping identifies causal variants for RA and T1D in *DNASE1L3, SIRPG, MEG3, TNFAIP3* and *CD28/CTLA4* loci

**DOI:** 10.1101/151423

**Authors:** Harm-Jan Westra, Marta Martinez Bonet, Suna Onengut, Annette Lee, Yang Luo, Nick Teslovich, Jane Worthington, Javier Martin, Tom Huizinga, Lars Klareskog, Solbritt Rantapaa-Dahlqvist, Wei-Min Chen, Aaron Quinlan, John A. Todd, Steve Eyre, Peter A. Nigrovic, Peter K. Gregersen, Stephen S Rich, Soumya Raychaudhuri

## Abstract

We fine-mapped 76 rheumatoid arthritis (RA) and type 1 diabetes (T1D) loci outside of the MHC. After sequencing 799 1kb regulatory (H3K4me3) regions within these loci in 568 individuals, we observed accurate imputation for 89% of common variants. We fine-mapped^1,2^ these loci in RA (11,475 cases, 15,870 controls)^3^, T1D (9,334 cases and 11,111 controls) ^4^ and combined datasets. We reduced the number of potential causal variants to ≤5 in 8 RA and 11 T1D loci. We identified causal missense variants in five loci (*DNASE1L3*, *SIRPG*, *PTPN22*, *SH2B3* and *TYK2)* and likely causal non-coding variants in six loci (*MEG3, TNFAIP3, CD28*/*CTLA4*, *ANKRD55*, *IL2RA*, *REL*/*PUS10*). Functional analysis confirmed allele specific binding and differential enhancer activity for three variants: the *CD28/CTLA4* rs117701653 SNP, the *TNFAIP3* rs35926684 indel, and the *MEG3* rs34552516 indel. This study demonstrates the potential for dense genotyping and imputation to pinpoint missense and non-coding causal alleles.

---

RA is an autoimmune disease in which chronic inflammation leads to joint destruction, which is associated with autoantibodies to citrullinated proteins in the majority of cases^5^. T1D arises through autoimmune destruction of pancreatic beta-cells, leading to complete loss of insulin production. Autoantibodies in T1D include those reactive to proinsulin^6^ and glutamic acid decarboxylase^7^. Genome wide association studies (GWAS) have identified over 101 RA loci^3,8^ and 53 T1D loci^4^. In order to define causal variants, fine-mapping has now been successfully applied to complex disease loci including inflammatory bowel disease^9^, type 2 diabetes^1,^^10^, coronary artery disease^1^, Graves disease^1^, and multiple sclerosis^11^. Since causal variants for both RA and T1D diseases overlap functional elements in CD4+ T cells^12^, we fine-mapped autosomal non-MHC loci for both diseases together.

We used ImmunoChip data for RA (11,475 cases, 15,870 controls)^3^, and T1D (9,334 cases and 11,111 controls; **Supplementary Table 1**)^4^. This platform contains dense coverage of single nucleotide polymorphisms (SNP) in selected autoimmune disease loci, enabling accurate imputation. Among these loci, 46 and 49 non-MHC autosomal loci have known significant associations for RA and T1D, respectively. Since RA and T1D share 19 loci, we examined 76 unique ImmunoChip loci in total (**Supplementary Table 2**).

We used three high-quality reference panels and selected the imputation strategy that maximizes coverage and accuracy for common variants (minor allele frequency; MAF>1%): 1) the Haplotype Reference Consortium (HRC, v1.1) reference panel (consisting of 64,976 haplotypes from 20 independent sequencing studies^13^), 2) the 1000 genomes (1KG, 3v5) European subpopulation (EUR) and 3) the 1KG cosmopolitan panel (COSMO)^14^. To evaluate accuracy of each strategy, we sequenced 568 individuals genotyped on ImmunoChip, targeting 799 1,000 bp regions centered around H3K4me3 peaks in ImmunoChip regions (**Online Methods**). From this data, we called 1,862 variants (MAF>1%; **Supplementary Figure 1**; **Supplementary Table 3**), which we compared to imputed genotypes. EUR and COSMO provided higher accuracy, compared to the HRC (89% vs 84% of variants with r^2^_g_>0.5; **Figure 1A&B**, **Supplementary Tables 4-5**). Imputation with COSMO obtained 1.8% higher coverage for variants with high quality (INFO>0.3) variants than with EUR (**Supplementary Table 6**). The difference between COSMO and HRC was partially due to the inclusion of insertion/deletion variants (indels) in COSMO (**Supplementary Figure 2 and 3**). INFO-scores were consistent with imputation accuracy (**Supplementary Figure 4**). We therefore opted to use COSMO to impute genotypes.

**Figure 1.**
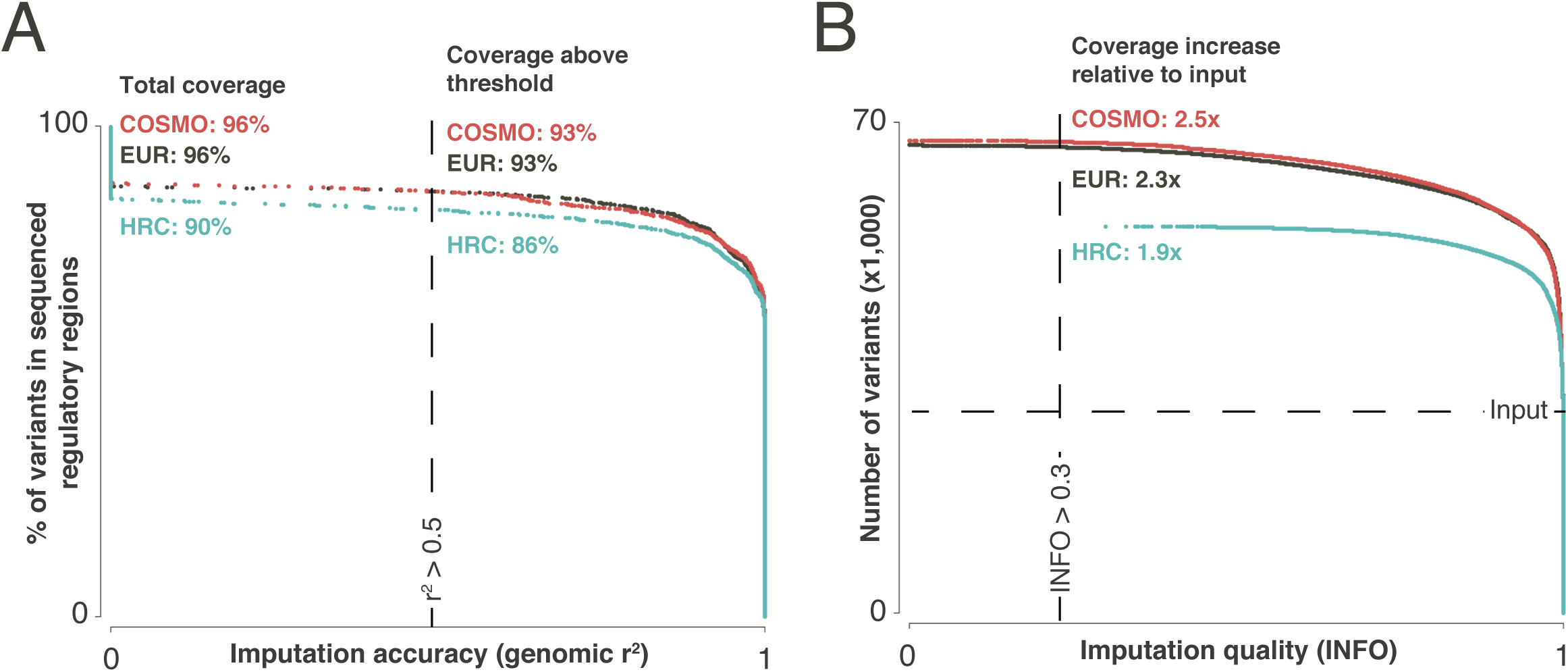
We imputed our datasets with different reference panels: the European subpopulation of 1000 genomes (EUR), full 1000 genomes (COSMO), and the Haplotype Reference Consortium (HRC). A) We sequenced 799 1kb regions in 568 individuals with ImmunoChip genotypes, and called 1,862 common (MAF>1%) variants. Imputation with COSMO and EUR recovers 96% of these variants, while HRC imputation recovers 90%. We calculated imputation accuracy by correlating imputed genotypes with genotypes called from the sequencing experiment (genomic r^2^). At r^2^>0.5, COSMO and EUR recover 93% of variants, while HRC recovers 86%. B) Imputation quality scores (INFO) for each reference panel in the RA dataset. COSMO shows highest increase in number of variants (MAF>1%) after imputation (2.5x; INFO>0.3) compared to EUR (2.3x) and HRC (1.9x).

Notably, even this best performing strategy had incomplete variant coverage: 4% of common variants in the targeted sequencing experiment were missed altogether, of which 75% were indels and multi-allelic variants (**Supplementary Figure 5**).

We focused our analysis on a subset of the loci with a tractable number of putative causal variants within our data set. First, we calculated association statistics for 64,430 and 66,115 imputed and genotyped variants for RA and T1D (MAF>1%, INFO>0.3; Hardy-Weinberg p>10^-5^) in the 76 loci. We observed association in 20 and 36 loci, for RA and T1D (p<7.5x10^-7^=0.05/66,115 tests; **Supplementary Table 7 and 8**). For 50% (=10/20) of RA and 72% (=26/36) of T1D loci, the most significant variant was in linkage disequilibrium (LD; r^2^>0.8) with the most significant previously published variant (**Supplementary Table 7 and 8**). RA and T1D variant effect sizes were positively correlated in 64% of the tested loci (**Online methods**, **Supplementary Table 9, Supplementary Figure 6**) suggesting shared signals. We therefore analyzed a combined dataset with 20,787 (RA or T1D) cases and 18,616 unique controls (**Online methods**). We restricted our analysis to 28 loci with sufficient statistical signal to warrant fine-mapping in the combined dataset (p<7.5x10^-7^). In the combined dataset, the strongest associated variant was in strong LD with the strongest associated variant in either RA or T1D in 69% of significant loci (r^2^>0.8; **Supplementary Table 10 and 11**). To prioritize loci with causal variation that we might be able to pinpoint, we created 90% credible sets using an approximate Bayesian approach^1,2^ and limited subsequent analysis to the 10 (RA), 15 (T1D) and 11 (combined) loci having ≤10 variants in the 90% credible sets (**Figure 2A&B; Supplementary Table 12**). Within the significant loci, we observed a striking 18.3-fold posterior probability enrichment for missense variants.

**Figure 2.**
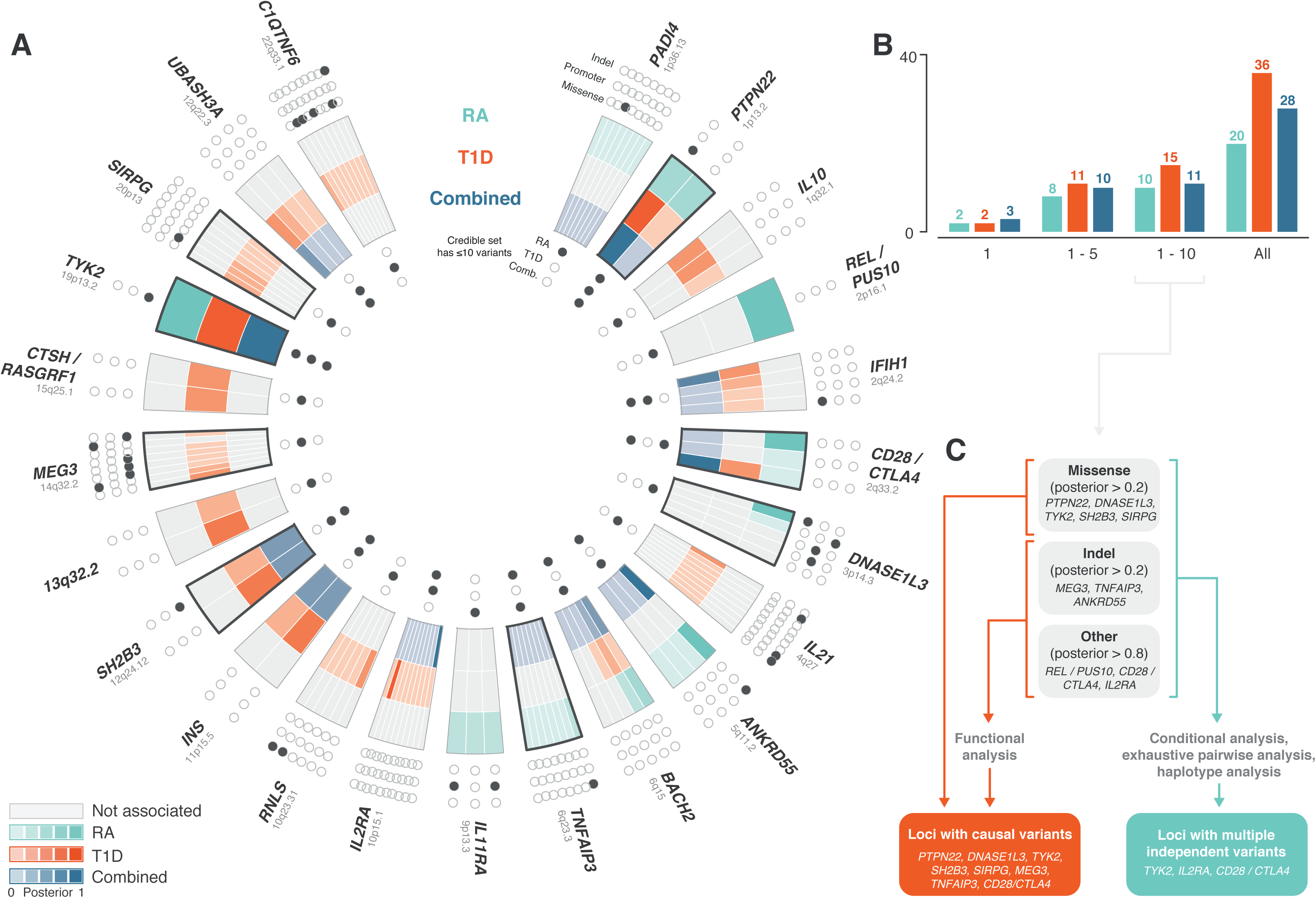
We used the approximate Bayesian factor to determine 90% credible sets within significant loci. A) A number of loci have a shared signal between diseases. Inner ring of dots indicates whether locus has ≤10 variants in credible set and has a significant association signal, and is open otherwise. Middle ring shows variants in each credible set. Highlighted segments indicate loci with causal variant. Color intensity indicates posterior probability and grey when not significant. Outer ring shows indel, promoter and missense coding annotation for each variant in credible set. B) We are able to narrow down the list of causal variants 5 in 8 out of 20 significant RA loci, and 11 out of 36 significant T1D loci. For both diseases, we find two loci that are explained by a single variant. C) From the credible sets, we defined groups of interesting loci, based on the presence of a high posterior missense variant (>0.2), indel (>0.2) or SNP (>0.8). We applied several follow-up analyses to these loci, including conditional analysis, exhaustive pairwise and haplotype analysis when a secondary signal was present, and functional analysis (EMSA) for non-coding loci.

We identified those alleles as likely causal if they had strong statistical genetic evidence and evidence of altered function (**Table 1**). To define strong candidate alleles, we defined three overlapping categories of promising loci: loci with 1) a single variant with a very high posterior probability (>0.8, *DNASE1L3, PTPN22, TYK2, CTLA4/CD28, REL/PUS10, IL2RA*), 2) a single missense variant with a modest posterior probability (>0.2, *DNASE1L3, PTPN22, SH2B3, TYK2, SIRPG*), or 3) a single non-coding indel with a modest posterior probability (>0.2, *TNFAIP3*, *MEG3, ANKRD55;* **Figure 2C; Supplementary Table 12**). We applied more modest thresholds to missense variants and indels, since they are *a priori* more likely to be functional. We considered high probability non-coding variants causal only if they met stringent additional criteria criteria suggesting function: 1) they occurred in a region with evidence of enhancer activity and 2) they demonstrated clear allele specific binding and enhancer function in vitro in both EMSA and luciferase assays.

**Table 1.**
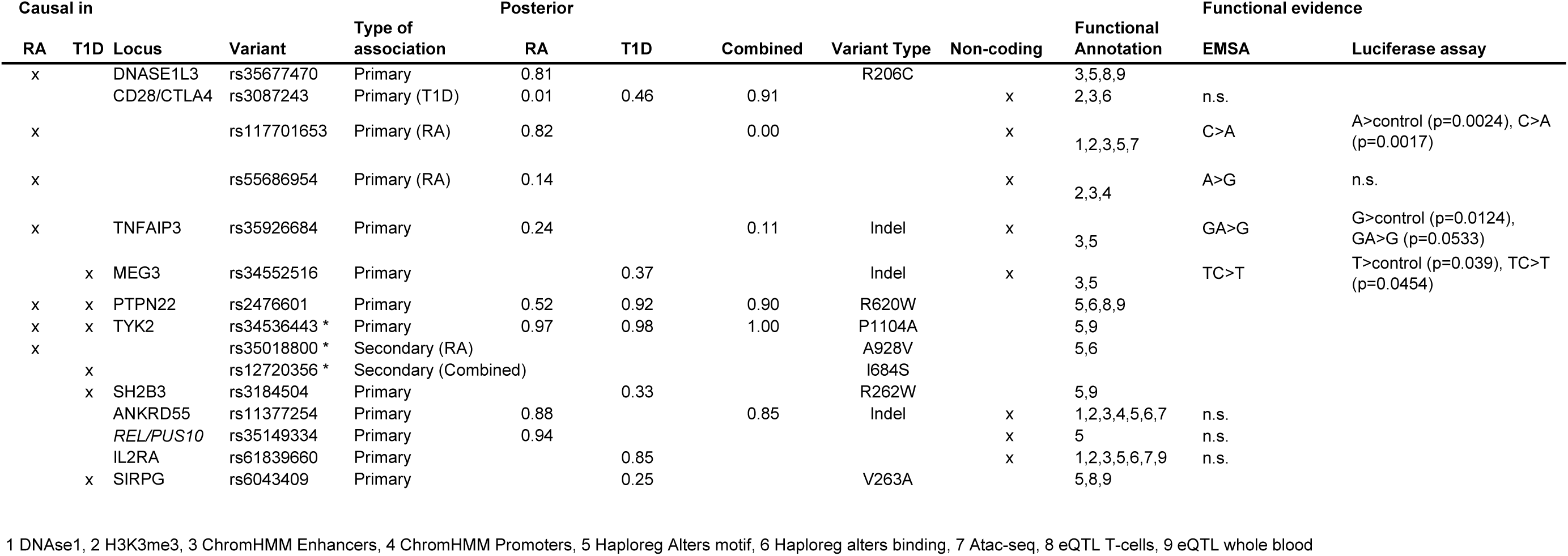
Overview of causal variants in selected loci. * identified using lower MAF threshold of 0.005. Greyed out posteriors are not significant in the primary association analysis. Functional annotations: 1 DNAse1, 2 H3K3me3, 3 ChromHMM Enhancers, 4 ChromHMM Promoters, 5 Haploreg Alters motif, 6 Haploreg alters binding, 7 ATAC-seq, 8 eQTL T cells, 9 eQTL whole blood. n.s.: non-specific binding.

We identified missense variants at *DNASE1L3* and *SIRPG*. We also identified causal missense variants at *PTPN22, SH2B3,* and *TYK2,* which are well described in the literature^4,15–17^ (**Supplementary Note and Supplementary Figures 7-9)**. Their identification suggests the sensitivity of our approach is high.

The 3p14 *DNASE1L3* locus, strongly associated with RA, but not T1D (p>0.02, **Supplementary Figure 10**), had a missense variant with high posterior probability. The previously reported^3^ lead SNP rs35677470 was included as one of the 5 variants within the 90% credible set of causal variants (p=1.7x10^−8^; posterior=0.81; **Supplementary Table 12**), and encodes a R206C change in the *DNASE1L3* protein product. After conditioning on R206C, we observed no evidence of independent risk variants (p>5x10^-4^; **Supplementary Table 13**). R206C has been implicated with systemic sclerosis^18^ and other loss of function mutations in *DNASE1L3* have been reported in familial forms of systemic lupus erythematosus^19^. R206C is a loss of function allele that abolishes the protein product’s nuclease activity^20^.

Within the *SIRPG* locus, we identified a missense variant with high posterior (rs6043409; p=3.94x10^-10^; posterior=0.25), causing a V263A substitution in the SIRPG gene product (**Supplementary Figure 11; Supplementary Table 12**). Conditional analysis using rs6043409 obviated the association signal in the locus (p>2x10^-3^). Since the consequence of V263A substitution on SIRPG function has yet to be described, we nominate it as a causal variant with caution.

Next, we focus on non-coding likely causal variants. We identified non-coding allele specific function in *CTLA4/CD28, TNFAIP3,* and *MEG3* using EMSA and luciferase assays in regions with evidence of CD4+ T cell enhancer function (**Table 1**). Loci having candidate variants with high posterior probabilities, but without evidence of allelic function, are presented in the **Supplementary Note and Supplementary Figures 12-14.**

The *CD28/CTLA4* locus has previously been shown to have shared association signals for RA and T1D^21^ and variant effect sizes between diseases are highly correlated in our analysis (Spearman’s rank r=0.9; **Supplementary Table 9**). In the combined analysis, we observed a single credible variant (rs3087243; p=1.4x10^-16^; posterior=0.91) near *CTLA4*. That same variant has the largest posterior probability in T1D (p=1. 7x10^-15^; posterior=0.46; **Figure 3A; Supplementary Figure 15A; Supplementary Table 12**), but not in RA (p=1.6x10^-8^; posterior=0.01). In contrast, in RA the rs117701653 variant carries high posterior probability (p=1.3x10^-10^; posterior=0.82); it is located closer to *CD28* and is not linked to rs3087243 (r^2^=0.03). Conditioning on rs3087243, we observed an independent effect at rs117701653 in RA (p=1.8x10^−8^), (**Figure 3A**; **Supplementary Table 13**). To confirm the two independent effects, we tested all pairs of SNPs exhaustively and observed that the rs3087243+rs117701653 pair demonstrates the most significant association of all SNP pairs in RA (**Figure 3B, Supplementary Figure 15B**). Haplotype analysis confirmed the independent protective effects of the rs3087243 A allele and of the rs117701653 C allele in both RA and T1D (**Figure 3C**), suggesting that rs117701653 may contribute to risk similarly in T1D (p=0.03 in conditional haplotype analysis).

**Figure 3.**
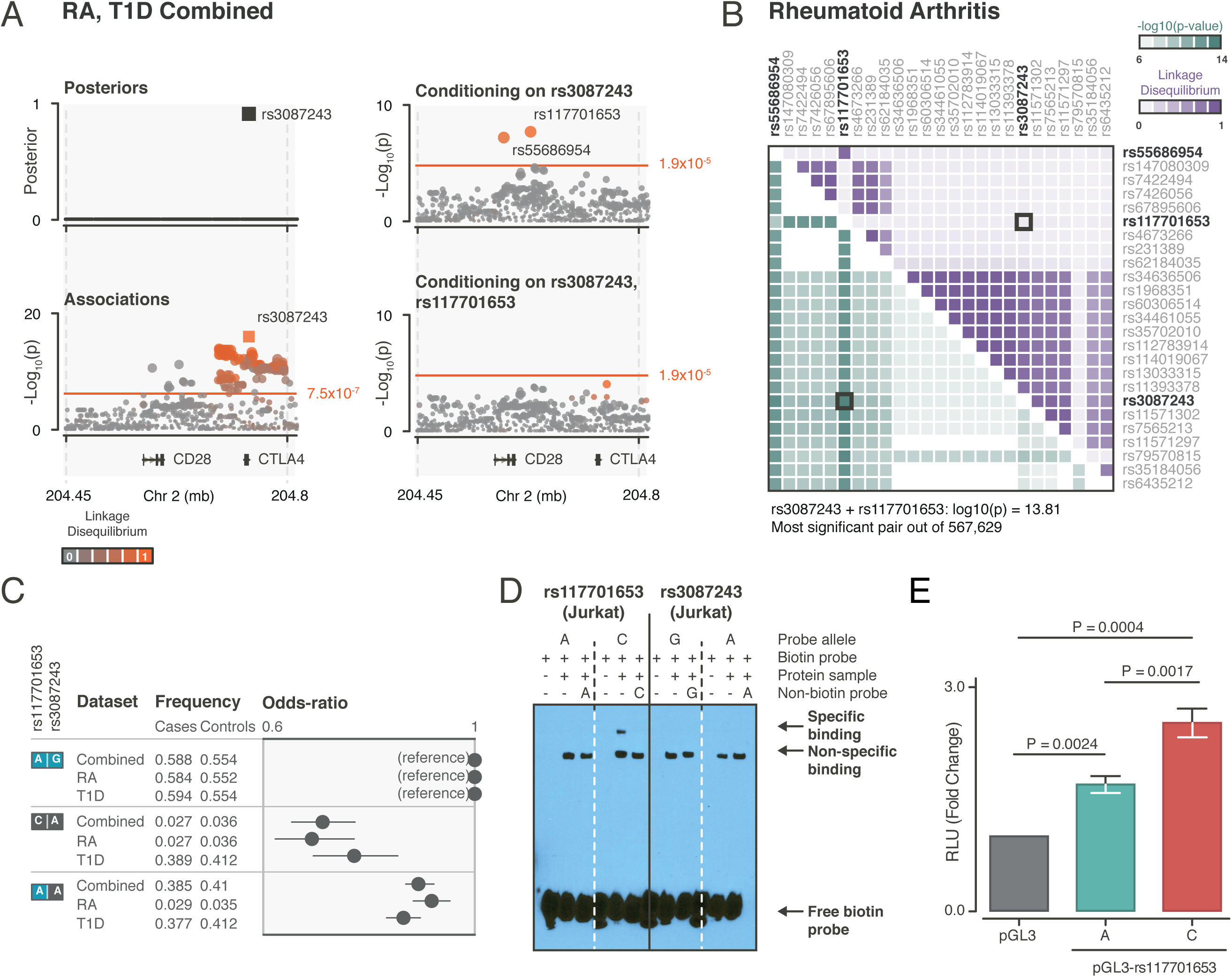
Analysis in the *CD28/CTLA4* locus. A) The regional association plot for the combined analysis shows a single variant (rs3087243), near *CTLA4*, in the credible set. Conditioning on rs30872043 reveals rs117701653 as an independent association. Color indicates LD with top-associated variant. Square indicates presence in credible set. B) Exhaustive pairwise analysis shows that rs3087243+rsrs117701653 pair has strongest association. Green color indicates-Log10(pairwise p-value), purple color indicates pairwise LD. C) Haplotype analysis using rs30872043 andrs11701653, using the AG haplotype as reference. The C allele of rs117701653 shows largest decrease in risk in RA, and the A of rs30872043 in T1D. D) EMSA using probes for rs117701653 and rs3087243 as a functional follow-up in Jurkat T cells. We observe an extra band in the lane with protein sample and biotin probe for the C-allele that is not observed for the other probes. The band disappears when adding non-labeled probe, suggesting competition between labeled and non-labeled probe for binding protein. E) Luciferase assay for rs117701653 using pGL3 plasmids in Jurkat T cells. We calculated relative luciferase activity units (RLU) using the activity of the empty plasmid (pGL3) as reference, and observed significant increase in luciferase activity for the A allele, and a further significant increase in luciferase activity for the C allele, which verifies that both alleles affect protein binding, albeit likely with different affinities.

We observed that both rs117701653 and rs3087243 may have regulatory function since they overlap H3K4me3 peaks in immune cell types, and disrupt protein binding motifs (**Supplementary Table 14 and 15**). Since regulatory regions can be context specific, we stimulated CD4+ T cells using CD3/CD28 beads, and measured chromatin accessibility using ATAC-seq before and after stimulation. We observed ATAC-seq peak overlap for rs117701653 only after stimulation (**Supplementary Table 16**), suggesting that rs117701653 may function specifically in stimulated cells. We note that while we did not observe linkage to eQTL in whole blood or T cells (**Supplementary Table 17**), rs3087243 did show a significant eQTL on *CTLA4* in testis^22^.

We demonstrated allele specific binding for rs117701653 but not rs3087243 using EMSA with Jurkat T cells (**Figure 3D**). The rs117701653 C allele showed higher specific binding affinity compared to the A allele (**Supplementary Figure 15C**). We also observed higher luciferase expression induced by the C allele compared to the A allele (p=0.0017; **Figure 3E**), suggesting allele specific enhancer activity. The binding is lineage specific: it was absent with THP-1 monocytic cells (**Supplementary Figure 15C**). As a relevant negative control, we also tested the second variant in the RA credible set (rs55686954; posterior=0.14), which showed no evidence of allele specific enhancer function (**Supplementary Figure 15C&D**). Published promoter capture Hi-C assays^23^ show local genomic contacts between the region harboring the rs117701653 SNP, the *CTLA4* promoter, and a region downstream of *RAPH1* (**Supplementary Figure 16**), indicating this allele might be regulating *CTLA4* or *RAPH1* despite proximity to *CD28.*

The *TNFAIP3* locus is associated with multiple autoimmune diseases^24–30^, including RA, but not T1D (p>2.3x10^-4^). We observed that the indel rs35926684 carries the highest posterior probability (p=6.8x10^-12^; posterior=0.24; **Figure 4A**; **Supplementary Table 12; Supplementary Figure 17A**) of 7 variants in the RA credible set. Conditional analysis revealed an independent association at rs58721818 (p=3.6x10^-5^; LD R^2^=0.05 with rs35926684; **Figure 4A; Supplementary Table 13**). A previous study^3^ identified rs6920220 (linked to rs35926684; r^2^=0.88) as the primary signal and secondary signals from rs5029937 (linked to rs58721818; r^2^=0.84) and rs13207033. Exhaustive pairwise analysis demonstrated comparable association for rs35926684+rs58721818 pair (-log10(p)=13.94) and the most strongly associated rs6920220+rs58721818 pair (-log10(p)=14.14; **Figure 4B; Supplementary Figure 17B**). Haplotypes having the rs35926684 G allele increased risk for RA, even in absence of the highly linked rs6920220 A risk allele (i.e. GGGC vs GAGC; **Figure 4C**), although this effect was only suggestive in conditional haplotype analysis (p=0.14).

**Figure 4.**
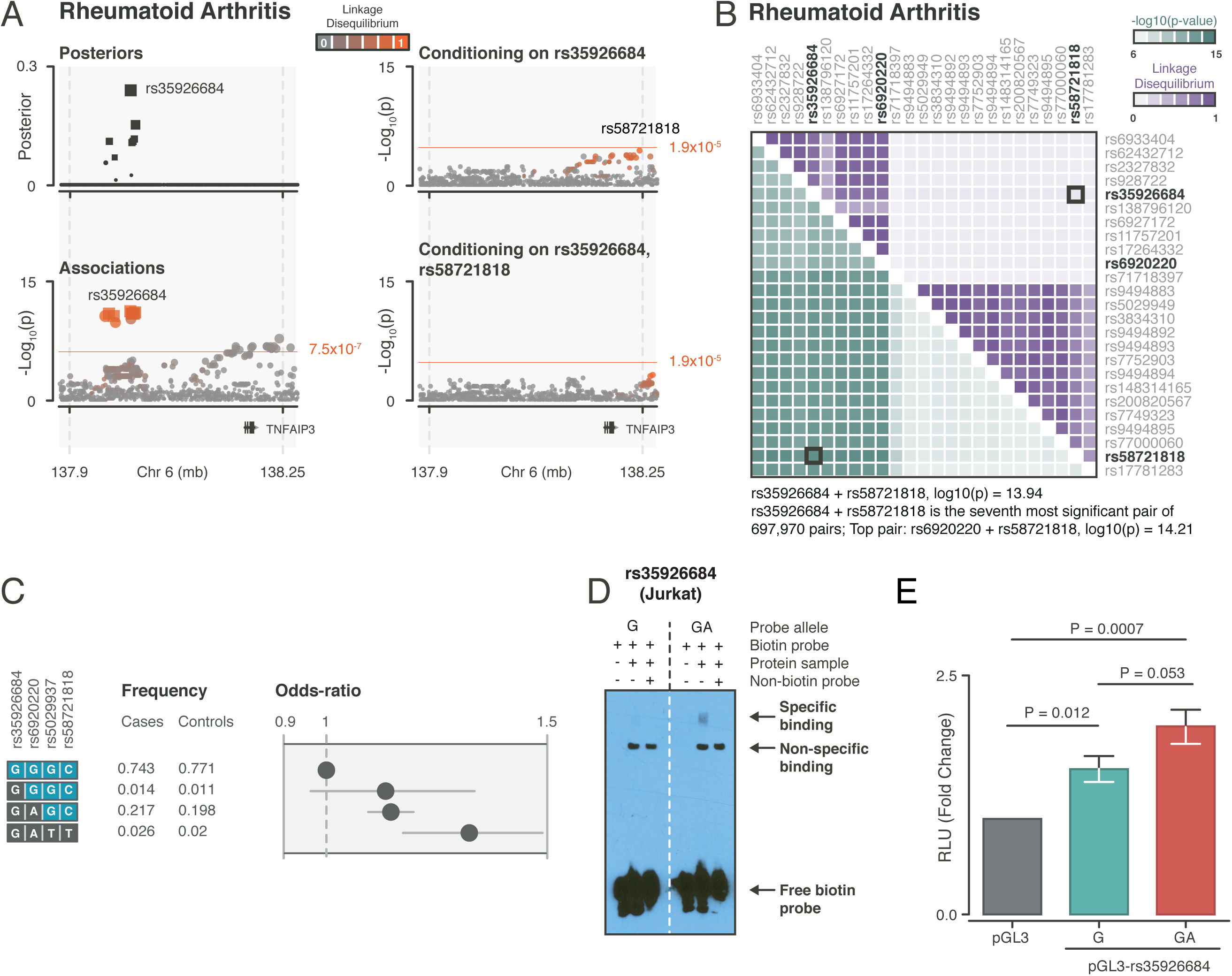
Analysis in the *TNFAIP3* locus. A) The variant with the strongest posterior in this locus is rs35926684, a G/GA indel, associated with RA. Conditional on rs35926684, we observe a significant secondary association with rs58721818. B) Exhaustive pairwise association analysis in RA indicates that there are 6 pairs with a lower p-value than rs35926684+rs58721818, although the top-associated pair (rs69220220+rs58721818) has an equivalent p-value (-log_10_(p)=13.94 vs 14.21). C) Haplotype analysis with rs35926684+rs58721818, and previously reported variants rs6920220 and rs5029937 shows that rs35926684 and previously reported top variant rs6920220 are often located on the same haplotype (GAGC), although a rare haplotype exists with only the alternative allele of rs35926684, which causes a similar increase in risk, although with larger standard error. D) EMSA analysis using a G and GA probe for rs35926684. We observe an extra band in the lane with protein sample and biotin probe for the GA-allele that is not observed for the other probes. The band disappears when adding non-labeled probe, suggesting competition between labeled and non-labeled probe for binding protein. E) Luciferase assay for rs35926684 shows that both G and GA consequently alter luciferase expression.

The rs35926684 indel alters more binding motifs, and overlaps more enhancer marks in immune related cell types, compared to rs6920220 (**Supplementary Table 14 and 15)**. Neither rs35926684 nor rs6920220 overlapped open chromatin region in our ATAC-seq time course (**Supplementary Table 16**), nor were linked with eQTLs in whole blood or T-cells (**Supplementary Table 17**).

EMSA with Jurkat cells demonstrated specific binding for rs35926684 (**Figure 4D**). Dose titration of the probe demonstrated specific binding for both G and GA allele, but stronger GA binding (**Supplementary Figure 17C**). Luciferase assays also demonstrated increased enhancer activity with the GA-allele compared to both the empty vector (p=7x10^-4^) and the G allele (p=0.053, **Figure 4E**). We did not observe specific binding with THP-1 cells, indicating cell type specificity (**Supplementary Figure 17C**). As a relevant negative control, we observed no allele specific binding for rs6920220 (**Supplementary Figure 17C**). Interestingly, in previously published promoter capture Hi-C data, the rs35926684 region contacts the *TNFAIP3* promoter^31^ as well as the *IL22RA* and *IFNGR1* promoters (**Supplementary Figure 16**)^23^, suggesting genes with immune function may be influenced by this RA risk allele.

*MEG3* is a non-coding RNA tumor suppressor gene whose transcript binds to p53^32^. In T1D, this locus has previously been described as an imprinted region, with greater risk carried by paternally inherited alleles^33^. We observed two variants in the credible set for T1D in this locus: the rs34552516 indel (p=1.1x10^-9^; posterior=0.37) and the rs56994090 intronic variant (p=7.3x10^-10^; posterior=0.54, **Figure 5A; Supplementary Figure 18A; Supplementary Table 12**). The locus shows no association with RA (p>0.04). Conditioning on rs34552516, we observed no evidence of additional independent effects (p>0.04; **Supplementary Table 13**). Both variants overlap DNAse-I sensitive, H3K4me1, and H3K4me3 regions in multiple immune cell types (**Supplementary Table 14**), but do not overlap open chromatin regions in our ATAC-seq experiment (**Supplementary Table 16**).

**Figure 5.**
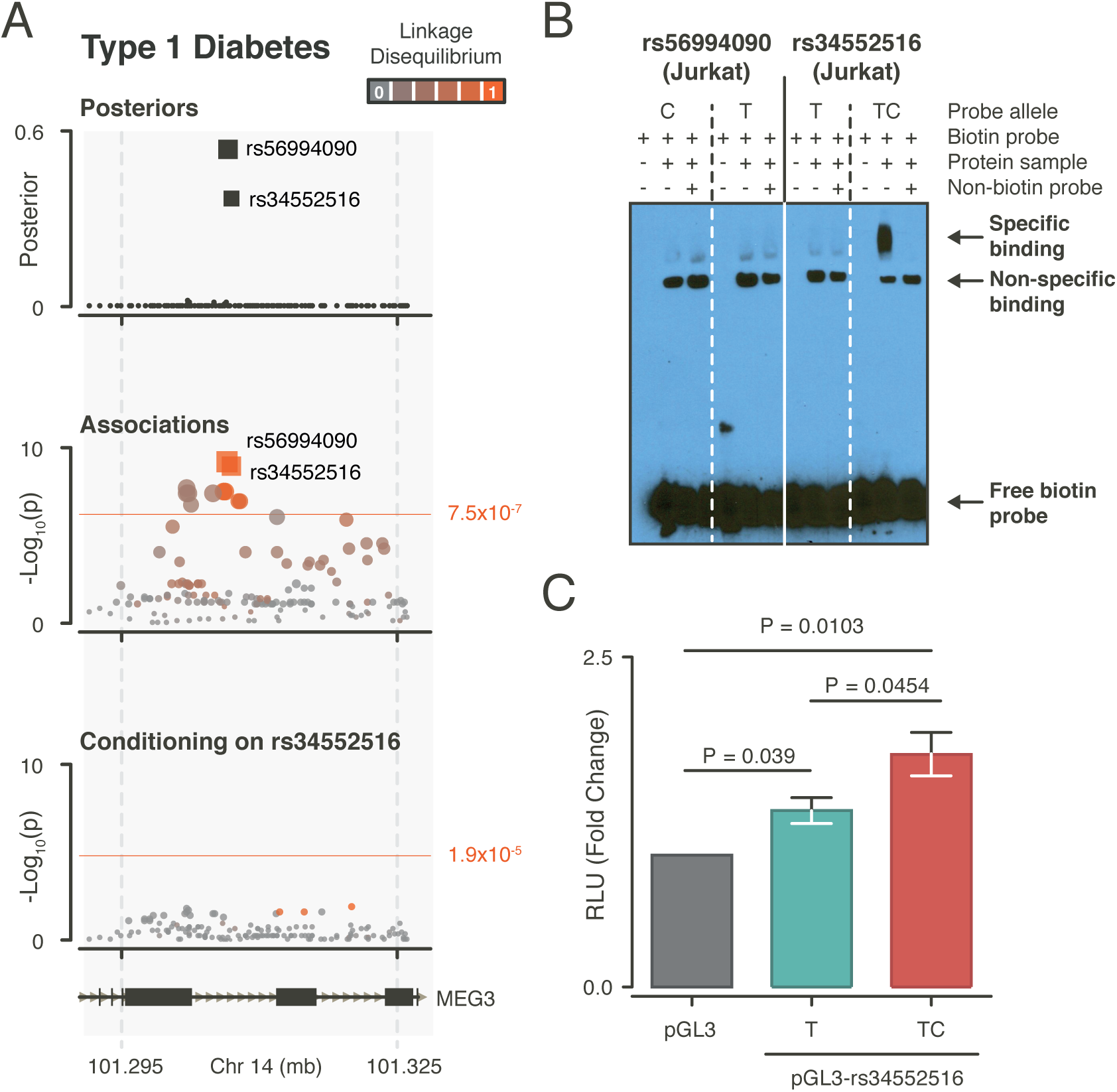
Analysis in the *MEG3* locus. A) Region plot for the *MEG3* locus in T1D. We observe two variants in the credible set (rs56994090 and indel rs34552516; indicated by squares). We did not observe secondary signals when conditioning on rs56994090. B) EMSA analysis for rs354552516 and rs56994090. Left: the TC allele of rs34552516 shows a band that disappears when adding non-labeled TC probe as competitor, suggesting specific binding. C) Consequently, a luciferase assay for rs34552516 shows an increase of luciferase activity for the TC allele relative to the T allele and empty vector.

EMSA with Jurkat cells showed protein binding specific to the TC allele of rs34552516 (**Figure 5B**), and the rs34552516 TC allele showed a significant increase in luciferase activity compared to empty vector (p=0.01) and the T allele (p<0.05; **Figure 5C**). We observed no specific binding with THP-1 cells (**Supplementary Figure 18B)**, indicating lineage specificity (**Figure 5B**). As a relevant negative control, we did not observe allele specific binding for rs56994090. The region harboring rs34552516 in promoter-capture Hi-C data^23^ showed contacts, including the promoter of *DIO3* and *RP11-1029J19* (**Supplementary Figure 16**), indicating that this risk allele may affect interaction with multiple downstream genes.

In this study, we identified three non-coding causal alleles with high posterior probability based on association data, and evidence of allele specific binding or enhancer function (**Table 1**). We observed in targeted sequencing that a proportion of causal variants might be missed by any imputation strategy, particularly indels or multiallelic variants. We therefore recognize that attempting to fine-map other loci may be more successful once more complete reference panels based on whole genome sequencing data become available, such as through the TopMed initiative (https://www.nhlbiwgs.org/).

Notably, the non-coding causal variants that we identified did not overlap with eQTL in either whole blood or T cells (**Supplementary Table 17**). Therefore, to elucidate the mechanisms underlying these variants, studies will be required to identify the precise protein complexes that bind these enhancers, and the downstream functions of those complexes.

We also identified other non-coding variants with high posterior probabilities that could feasibly be pursued for validation, but did not demonstrate clear evidence of allele-specific function in our assays. Other more sensitive assays, or application of assays in other non-CD4+ T cell-types might ultimately be able to confirm the function of these alleles too.

## Data Availability

Summary statistics for all variants will be made available upon acceptance. Genotype data is previously published^3,4^ and is available from RACI and

T1DGCC upon request. ATAC seq data will be deposited upon acceptance of this manuscript to GEO.

Bios eQTL browser: http://genenetwork.nl/biosqtlbrowser/, Roadmap epigenomics datasets: http://www.roadmapepigenomics.org/, ChromHMM enhancers and promotors: http://egg2.wustl.edu/roadmap/web_portal/, 1000 genomes reference panel: http://bochet.gcc.biostat.washington.edu/beagle/1000_Genomes_phase3_v5a/, Haplotype Reference Consortium panel: http://www.haplotype-reference-consortium.org/

## Code Availability

Associated computer code for this manuscript can be found at the following GitHub repositories: https://github.com/immunogenomics/harmjan/tree/master/FinemappingPaper and https://github.com/immunogenomics/harmjan/tree/master/FinemappingTools

## Author Contributions

**Analysis**: H-J.W., Y.L., S.R.; **Functional Assays**: M.M.B., P.A.N.

**Study Design**: H-J.W., P.A.N., S.R.; **Data Acquisition**: S.O., A.L., N.T., J.W., J.M., T.H., L.K., S.R-D., W-M. C., A.Q., J.A.T., P.K.G., S.S.R., S.R.; **Writing and editing manuscript**: H-J.W., M.M.B., Y.L., J.A.T., P.A.N., P.K.G, S.S.R., S.R

## Acknowledgements

This work is supported in part by funding from the National Institutes of Health (U01GM092691, UH2AR067677, 1U01HG009088, and 1R01AR063759 (SR)), and the Doris Duke Charitable Foundation Grant #2013097. This work is part of the research program Rubicon ALW with project number #825.14.019 (HJW), which is (partly) financed by the Netherlands Organization for Scientific Research (NWO). Further support was provided by the Wellcome Trust [107212/Z/15/Z] and JDRF [5-SRA-2015-130-A-N] to the Diabetes and Inflammation Laboratory and by the Wellcome Trust [203141/Z/16/Z] to the Wellcome Trust Centre for Human Genetics. PKG was supported in part by the Feinstein Institute and a generous gift from Eileen Ludwig Greenland (PKG). PAN is supported by a Rheumatology Research Foundation Disease Targeted Research Grant, NIH P30 AR070253, and the Fundación Bechara.

## Online Methods

### Patient collections

We used genotyping data from samples that were collected on the ImmunoChip platform, which were obtained with informed consent and described in previous publications (**Supplementary Table 1**)^3,4^. In summary, for RA, we used ImmunoChip data for 11,475 cases and 15,870 controls, collected by six different cohorts (UK, Swedish EIRA, United States, Dutch, Swedish UMEA, and Spanish)^3^. For T1D, we used ImmunoChip data for 12,241 cases and 14,636 controls divided over two different cohorts, that have been described earlier^4^: the T1DGC family collection (T1D EUR) and the UK GRID, British 1958 Birth cohort and UK Blood Service collection (T1D UK). In order to include trios from the T1D EUR cohort in a case-control analysis, we generated pseudocontrol pairs for each affected individual using the untransmitted alleles from the parents of that individual. As a consequence, the final number of individuals for T1D was 9,334 cases, and 11,111 controls (including 1,661 pseudocontrols). Quality control on the genotypes was performed as described in the previously published studies. Additionally, we merged the genotype data for the different cohorts within T1D and RA using PLINK^34^, and converted genomic coordinates using the UCSC liftOver tool and the hg18ToHg19 chain file. Variants unable to liftOver were removed. We then replaced the variant identifiers using NCBI dbSNP build 138. Finally, we removed variants with a MAF < 0.5%.

### Imputation

In order to assess which imputation strategy was best suited for fine-mapping, we tested three reference panels: 1) The European subpopulation of 1000 genomes (N=503), 2) the cosmopolitan panel of 1000 genomes (N=2,504), and 3) the HRC v1.1 reference panel (N=32,611). Our approach used three steps (matching, imputation, and merging). First, we matched variants to each reference panel: we removed variants that were not present in the reference panel and aligned the strands of the remaining ImmunoChip genotypes. We excluded variants when alleles could not be matched, or in the case of C/G and A/T variants, when the minor allele was unequal. If we observed an unequal minor allele for such variants, and the reference panel and ImmunoChip MAF was >45%, we chose to flip the allele in the ImmunoChip data. For multi-allelic variants, we ensured that the allele encoding was identical relative to the reference panel variant. As a consequence of these steps, the input for each reference panel was slightly different (**Supplementary Table 4**). Second, we imputed genotypes into RA and T1D separately. We phased and imputed the 1000 genomes reference panels using Beagle v4.1 (version 22Apr16.1cf)^35^. In order to accommodate computational constraints of Beagle, we split the RA and T1D datasets into 30 batches, randomizing cases and controls between batches, while maintaining trio structure in the T1D dataset. Since the HRC v1.1 reference panel genotype data is not publicly available, we evaluated different imputation servers and settings for the T1D dataset, in order to determine their effects on imputation output. On the Sanger Institute imputation server (date of access: May 11, 2016), we used prephasing with either EAGLE^36^ or SHAPEIT^37^, followed by imputation with PBWT^38^. On the Michigan University server (date of access: July 5, 2016), we used prephasing with EAGLE^36^, followed by MiniMac^39^ imputation. Due to the constraints of the Michigan University imputation server website, we split the dataset into three batches, randomizing cases and controls while maintaining trios. For RA, we performed HRC imputation on the Sanger imputation server using EAGLE prephasing followed by PBWT imputation. Third, we merged the imputed dosages and probabilities from each batch (if any) for each imputation reference panel, and replaced the variant identifiers in the imputed output using NCBI dbSNP build 138. Before calculating association statistics, we replaced genotypes for variants genotyped on ImmunoChip with the original genotypes. Finally, we recalculated the imputation quality scores for each imputed variant in each dataset: for biallelic variants, we used the INFO score and Beagle v4.1 allelic-R^2^ for multi allelic variants.

### Targeted sequencing

In order to test the accuracy of imputation, we sequenced targeted regions in 864 individuals (160 T1D trios and 384 unrelated RA cases, of which 480 and 149 were on ImmunoChip, respectively). We used the Illumina MiSeq platform to generate 100bp paired-end reads. We sequenced 900 regions of 1,000bp around H3K4me3 peaks centers overlapping loci associated with either disease, since these loci are most likely to harbor causal variants^12^. We used BWA-mem^40^ (v0.7.12) to align reads to the hg19 reference genome. We tagged and removed duplicate reads using Picard MarkDuplicates. We then removed 101 regions where >50% of the samples had <20x coverage at >80% of sequenced bases, and removed 86 samples having <20x coverage at 90% of sequenced bases. We called genotypes using GATK version 3.4, following the recommended guidelines for using HaplotypeCaller^41^ in a joint genotype calling approach. We then set genotypes with <10x coverage and QUAL<30 to missing, and excluded variants with >5% missingness. We corrected for possible sample swaps and mismatched samples by correlating the called genotypes with ImmunoChip genotypes and removing samples that did not match any ImmunoChip sample (r<0.95), resulting in 568 final samples (including 439 for T1D, and 129 for RA). Finally, we selected variants with MAF>1%, resulting in 1,862 variants within the 76 RA and T1D associated regions.

### Merging imputed datasets

Prior to the association analysis, we merged the data for the RA and T1D dataset, imputed with the COSMO reference panel. Since these cohorts share controls, not necessarily with identical identifiers, we first identified individuals with high shared genetic background. For this purpose, we first generated a list of LD pruned variants from the ImmunoChip genotypes using PLINK^34^ (using -- indep-pairwise 1000 100 0.2). We then used this list to determine the genetic similarity (unified additive relationship; UAR)^42^ between each pair of samples across both datasets. We considered sample pairs with an UAR>0.2 genetically related and randomly selected one sample of the pair to be included from either the RA or the T1D dataset. We considered the remaining sample pairs unrelated. We finally merged genotypes and imputation probabilities from the selected samples, and recalculated the imputation INFO scores for the merged genotypes as described earlier.

### Association analysis framework

#### Fine-mapping and statistical analysis

Due to the sample size of the datasets in our study, we limited our association analysis to variants having an overall MAF>1%, a Hardy-Weinberg P-value (HWE-P)>10^-5^ in controls, and an overall INFO score>0.3. HWE-P was calculated using an exact test for biallelic variants, while for multi-allelic variants, Pearson’s chi-squared test was applied. We then split multi-allelic variants into multiple variants, creating a single variant for each alternate allele. To test each variant for association with disease, we used logistic regression, assuming a log-linear relation between the number of alternative alleles and case-control status. We then created a null model containing covariates in order to account of population differences. In the RA dataset, the null model included the first 10 principal components calculated over the genotype covariance matrix as described earlier^3^, and included an additional 5 covariates indicating the originating cohort. For T1D, we included 12 regional indicator variables in the null model as described earlier^4^, and an additional variable indicating the originating cohort. For each variant, we then fit an alternate model containing the genotypes. For the joint analysis, the null model included all covariates for the T1D and RA datasets and an additional covariate indicating whether the sample originated either from the RA or the T1D dataset. In order to account for imputation uncertainty, we recoded the imputation probabilities to a dosage value ranging between 0 and 2 (i.e. P(AB) + 2xP(BB)). Finally, we calculated the p-value for the association as the difference in deviance between the alternative and null models, which follows a chi-squared distribution with 1 degree of freedom. To determine the significance of the association we calculated a study-wide Bonferroni threshold using the maximum number of tests across datasets (p<7.5x10^-7^=0.05/66,115).

#### Definition of credible sets

To define the most likely causal variant for each locus, we calculated posterior p-values using the approximate Bayesian factor (ABF)^1,2^ under the assumption of a single causal variant per locus. Shortly, this framework assumes that the association effect sizes follow a N(0, V) distribution under H_0,_ with V being the standard error squared of the association. Under H_1_ the framework assumes a distribution following N(0,V+W), where W is (ln(1.5)/1.96)^2^, reflecting the prior of observing an odds ratio of 1.5. The ABF for an observed effect size ß is then calculated as the ratio of P(ß|H_0_)/P(ß|H_1_), effectively measuring the probability of observing the effect size under the H_0_ of no association over the H_A_ of observing an association. Using the sum of the ABF for all variants in the locus, we calculate the posterior for variant i as:

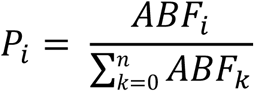

Following calculation of the posterior p-values, we created credible sets within each locus by sorting associations descending on the basis of their posterior p-values, and including associations such that the sum of their posteriors is >0.9.

#### Detecting secondary associations

In order to determine the presence of multiple independent effects, we performed a conditional analysis using logistic regression: for each locus with a significant association, we included the top-associated variant as a covariate in the null model, and repeated the association analysis for that locus.

For each locus with a significant secondary association, we then tested whether the observed pair of top-associated variants together provided the strongest pairwise association signal given the variants in the locus by performing an exhaustive pairwise analysis. Similarly to the normal logistic regression, the null model included the covariates for each dataset, while the alternate model included genotype dosages for both variants. The significance of the pairwise association was then calculated using the difference in deviance between the null and alternative models, following a chi-squared distribution with 2 degrees of freedom.

Finally, for loci with two or more independent associations, we assessed whether the risk alleles for the associated variants were located on the same haplotypes. For the independently associated variants, we derived haplotypes from the phased imputation output (e.g. 4 haplotypes for 2 independent variants), and assigned two haplotypes to each individual. We then removed all haplotypes with a frequency <1%, and removed all individuals that had any of the removed haplotypes from the analysis. By using the haplotype with the highest frequency as the reference haplotype, we assigned each individual to have either 0,1, or 2 copies of each alternative haplotype. We then used logistic regression to test each haplotype for association, assuming a log-linear relationship between the number of haplotype copies and disease status. To correct for population differences, our null model included covariates as described above.

### Functional annotation

#### eQTLs, H3K4me3 peaks, DNAse-I hypersensitive sites, enhancers and motifs

In order to provide functional annotation for the identified variants, we assessed overlap with eQTL, H3K4me3 peaks, DNAse-I hypersensitive sites, promoters and enhancers. We used eQTL from a large RNA-seq based eQTL meta-analysis using 2,116 whole blood samples^43^. Because many eQTL are cell type specific, and RA and T1D loci are enriched for enhancers in CD4+ T cells^12^, we also included a study assessing eQTL in CD4+ T cells using 461 individuals^44^. For each variant in a credible set, we first determined whether the variant was present in the eQTL summary statistics. We then selected the eQTL gene with the lowest eQTL p-value. For variants that were not present, we selected the eQTL snp with a linkage disequilibrium (LD) r^2^>0.8, using the European subpopulation in 1000 genomes as a reference panel. For eQTLs with equal LD, we selected the eQTL gene with the lowest P-value.

We downloaded annotations in narrowPeak format for H3K4me3 peaks, DNAseI peaks, and ChromHMM^45^ genome segmentations from the Roadmap epigenetics consortium^46^, consisting of 127 consolidated epigenomes from a large number of different cell types. We then grouped immune related cell types into an ‘immune’ group, and the remaining cell types in an ‘other’ group, resulting in two groups for DNAse-I and H3K4me3 annotations. We additionally used ChromHMM annotations created using 12 imputed epigenetic marks. Additional to the ‘immune’ and ‘other’ groups, we further grouped ChromHMM segments for enhancers (i.e. segments with an EnhA1, EnhA2, EnhW1, EnhW2, and enHAc annotation) and promoters (i.e. segments with PromP, PromBiv, PromU, PromD1 and PromD2 annotation), resulting in four annotation groups for ChromHMM annotations. Within each group, we subsequently determined the percentage of files in which we observed overlap between an annotation and variants within the credible sets. Finally, we determined whether candidate causal variants affected protein binding motifs or transcription factor binding sites using HaploReg^47^.

The number of cell types in each group was different between annotations, because not all annotations were present for all cell types. Numbers of files per annotation group can be found in **Supplementary Table 14**.

#### ATAC-seq timeseries

ATAC-seq is a method to measure chromatin accessibility using a small number of cells^48^. We here applied ATAC-seq to measure chromatin accessibility in a timeseries after stimulation. We used 30mL whole blood from a leukopak acquired from a healthy anonymous donor in a 20mL PBS solution. We then isolated PBMCs using Ficoll tubes and stored 500μl aliquots of 100x10^6^ cells in liquid nitrogen. Cells were subsequently thawed, and stained with anti-biotin microbeads for magnetic assisted cell sorting (MACS) to select CD4+ Tmem cells. Cells were resuspended and transferred to a 24 wells plate in 3ml aliquots of 6x10^6^ cells. Cells were stimulated using Dynabeads (Human T-Activator CD3/CD28 for T Cell Expansion and Activation; Life Technologies) in a 2 cells per bead ratio. Samples of 100,000 cells were taken at 0, 1, 2, 4, 8, 12, 24, and 48 hours after stimulation. Nucleosome isolation and ATAC-seq open chromatin sequencing was performed as described earlier^48^. Sequenced reads were mapped to the hg19 reference genome, using BWA-mem. Reads mapping to the mitochondrial genome, reads mapping to multiple genomic locations, and duplicate reads (labeled by Picard MarkDuplicates) were removed, and reads were shifted +4 and -5 bp for the reverse and forward strands respectively. Enrichment for open chromatin was determined by calling peaks using MACS v2^49^, using default settings.

### Electrophoretic mobility shift assay

#### Cell lines

Lymphocytic and monocytic cell lines, Jurkat and THP-1 respectively, were obtained from the ATCC (TIB-152 and TIB-202). Jurkat cells were grown in complete RPMI (RPMI-1640, Gibco, with 10% decomplemented-fetal bovine serum, penicillin and streptomycin) and THP-1 cells in complete RPMI supplemented with 2-mercaptoethanol to a final concentration of 0.05 mM. Both cell lines were grown in a 37°C incubator with 5% CO_2_.

#### Electrophoretic mobility shift assay (EMSA)

EMSA was performed using the LightShift Chemiluminiscent EMSA Kit (Thermo Scientific). Single stranded oligonucleotides corresponding to a 30-32 nucleotides fragment of the human genome with the SNP of interest in the middle were purchased from IDT (**Supplementary Table 18**). Single stranded oligonucleotides were biotinylated using the Biotin 3’End DNA Labeling Kit (Thermo Scientific) following manufacturer instructions. Double stranded oligonucleotides were generated by mixing together equal amounts of biotin-labeled (for probe) or unlabeled (for competitor) complementary oligonucleotides and incubating them 5 min at 95°C and then 1 hour at room temperature.

Nuclear extract from Jurkat and THP-1 cells was obtained using the NE-PER™ Nuclear and Cytoplasmic Extraction Reagents (Thermo Scientific) following manufacturer instructions. Protein extracts were dialyzed using a dialysis membrane with MWCO of 12-14 KDa (Spectrum Spectra) against 1 L of dialysis buffer (10 mM Tris pH 7.5, 50 mM KCl, 200 mM NaCl, 1 mM DTT, 1 mM PMSF and 10% glycerol) for 16 hours at 4°C with slow stirring. Protein inhibitor cocktail (Sigma-Aldrich) was added to a final concentration of 1.5x. Protein concentration was measured using the Pierce BCA Protein Assay Kit (Thermo Scientific) and adjusted to 4 μg/μl.

The standard binding reaction contained 2 μl of 10x Binding Buffer (100 mM Tris pH 7.5, 500 mM KCl and 10 mM DTT), 2.5% glycerol, 5 mM MgCl_2_, 0.05% NP-40, 50 ng Poly (dI:dC), 20 fmol biotin-labeled probe and 16 μg nuclear extract in a final volume of 20 μl. For competition experiments, a 200-fold molar excess (4 pmol) of unlabeled probe was added.

Binding reactions were incubated at room temperature for 30 min and loaded onto a 6 % polyacrylamide 0.5x TBE gel. After sample electrophoresis and transfer to a nylon membrane, transferred DNA was crosslinked for 10 min and the migration of the biotinylated probes and their complexes was detected by chemiluminescence followed by film exposure.

#### Luciferase reporter assay

The double stranded oligonucleotide containing the SNP of interest (obtained as described above) was cloned downstream the luciferase gene in the luciferase reporter vector pGL3 promoter (Promega). For that, unlabeled double stranded oligonucleotides containing the rs117701653, rs35926684 or rs34552516 were amplified with specific primers containing the BamHI restriction site obtained from IDT (**Supplementary Table 19**). The PCR was carried out in 50 μl reaction volume under the following program: 94°C 3 min; 10 cycles 94°C 30 sec, 60°C 40 sec, 68°C 30 sec; 15 cycles 94°C 30 sec, 60°C 40 sec, 68°C 30 sec; 72°C 10 min. Both PCR products and pGL3 promoter vector were digested with BamHI (New England Biolabs) for 1 h at 37°C and linearized vector was then dephosphorylated for 30 min at 37°C with the Quick Dephosphorylation kit (New England Biolabs). Digestion products were analyzed by electrophoresis in 1.2% agarose gels, and purified with QIAquick Gel Extraction Kit (Qiagen). Ligation of SNP containing fragments into the pGL3 promoter plasmid was performed in a ratio 1:50 (vector:insert) with T4 DNA ligase at 16°C overnight and transformed into JM109 competent cells (Promega). Plasmids from independent colonies were isolated using Wizard Plus SV minipreps DNA purification system (Promega) and sequenced using RV primer 4 (Promega) by Eton Bioscience. For each of the SNP, 3 colonies harboring the SNP-construct cloned “in sense” in the pGL3 promoter vector were selected for further plasmid isolation for transfection into Jurkat T cells.

Three independent transfection experiments for each construct were performed, each in duplicate. 0.6 x 10^4^ Jurkat cells in 0.1 ml of Opti-MEM (Gibco) were transfected with 0.8 μg of pGL3-promoter vectors, either without insert or with any of the six SNP-containing inserts, along with 0.2 μg of pRL-TK Renilla luciferase vector (Promega) using 1.5 μl of Lipofectamine LTX Reagent and 1 μl of PLUS Reagent (both from Invitrogen). After 16 hours of transfection, luciferase activity was measured using the Dual-Glo Luciferase assay system (Promega) following manufacturer instructions. Firefly luciferase activity was expressed as relative luciferase units (RLU) after correction for Renilla luciferase activity to adjust for transfection efficiency. Data were normalized to those cells transfected with empty pGL3-promoter vector. Results from the different clones were pooled together and expression levels compared by unpaired two-sided t-test.

## Supplementary Figures

**Supplementary Figure 1**

A) Out of the 902 sequenced regions, 799 had >20x coverage at 80% of sequenced bases in at least 50% of the samples. B) 87 samples had less than 20x coverage in at least 90% of the sequences bases. C) The 1,170 variants out of the 1,862 called variants that overlapped within 568 ImmunoChip genotyped individuals were highly correlated for both RA and T1D

**Supplementary Figure 2**

Top: Imputation quality (INFO) scores for the RA and T1D datasets, imputed with the European subpopulation of 1000 genomes (EUR), full 1000 genomes (COSMO), and the Haplotype Reference Consortium (HRC) reference panels. In T1D, HRC imputation was performed in three ways: using EAGLE (HRC / EAGLE) or SHAPEIT (HRC / SHAPEIT) for phasing on the Sanger Institute imputation server, or using EAGLE for phasing and imputation on the Michigan imputation server (HRC / EAGLE / MICHIGAN). From left to right: comparing variants with MAF>1%, comparing variants with MAF>1% but excluding indels, and comparing all variants. For MAF>1% variants, COSMO outperforms both EUR and HRC.

**Supplementary Figure 3**

Imputation accuracy (genomic r^2^) for the RA and T1D datasets, imputed with the European subpopulation of 1000 genomes (EUR), full 1000 genomes (COSMO), and the Haplotype Reference Consortium (HRC) reference panels. Genomic r2 was calculated by correlating imputed dosages with sequenced variants for the same individuals. In T1D, HRC imputation was performed in three ways: using EAGLE (HRC / EAGLE) or SHAPEIT (HRC / SHAPEIT) for phasing on the Sanger Institute imputation server, or using EAGLE for phasing and imputation on the Michigan imputation server (HRC / EAGLE / MICHIGAN). From left to right: comparing variants with MAF>1%, comparing variants with MAF>1% but excluding indels, and comparing all variants. In all cases, COSMO outperforms both EUR and HRC.

**Supplementary Figure 4**

We compared imputation quality (INFO score) with imputation accuracy (genomic r^2^) in the T1D dataset, and observed a strong correlation (r^2^=0.82).

**Supplementary Figure 5**

In the T1D dataset, 72 variants (MAF>1%) that were present in our gold standard genotype dataset were not present after imputation. A) The majority (69%) of these variants were indels and B) variants of low allele frequency (44% with MAF<5%). C) For those variants with a low MAF (MAF<1%), or with a low correlation with gold standard genotypes (r2<0.5), the majority (77%) were low frequency variants.

**Supplementary Figure 6**

In 66% of the 76 tested loci, the association statistics (Z-scores) between RA and T1D are positively correlated.

**Supplementary Figure 7**

Region plot for the *PTPN22* locus. The credible set consists two variants (indicated by squares): we observe two significant (p<7.5x10^-7^) associations in RA and T1D. These associations include rs2476601, which causes a R620W coding change in the PTPN22 protein and has a high posterior (0.78) in the combined analysis. No significant secondary signals are observed when conditioning on rs2476601. Color indicates LD between top associated variant.

**Supplementary Figure 8**

A) Region plot for the *TYK2* locus. Considering previous analysis in this region, we decreased the MAF threshold for this region to 0.5%. For each analysis, the credible set consists of a single variant, rs34536443, causing a P1104A change in TYK2. Conditional on P1104A, we observe a secondary association from rs35018800 in RA, causing a A928V change in TYK2. Further conditioning indicates a tertiary association from rs12720356 in the combined analysis, causing a I684S change in TYK2. Finally, conditional on these three coding variants, we observe a quaternary association from rs35074907. B) Top 25 SNPs as identified by pairwise exhaustive analysis. In RA and the combined analysis, rs34536443+rs35018800 is the top associated pair. In T1D, however, there are 138 pairs with a lower p-value, with rs35018800 + rs12720356 having the strongest association. C) Haplotype analysis using rs34536443, rs12720356, rs35018800, and rs35074907 using the GGAG haplotype as a reference. All haplotypes confer independent relative risk reduction, except for GGAA, which increases risk in T1D, relative to the reference haplotype.

**Supplementary Figure 9**

Region plot for the *SH2B3* locus. The credible set for T1D contains two variants, including rs3184504, causing a R262W change in SH2B3. Conditioning on rs3184504, we do not observe a secondary association.

**Supplementary Figure 10**

Region plot for the *DNASE1L3* locus. The credible set consists of two variants in RA, including rs35677470, causing a R206C coding change in the DNASE1L3 protein. No significant secondary signals are observed when conditioning on rs35677470.

**Supplementary Figure 11**

Region plot for the *SIRPG* locus. The credible set consists of seven variants in T1D, including rs6043409, causing a V263A coding change in the SIRPG protein. No significant secondary signals are observed when conditioning on rs6043409.

**Supplementary Figure 12**

Region plot for the *ANKRD55* locus. We did not observe a significant association for T1D, while for RA, the credible set contained two variants: rs11377254 and rs7731626 (indicated by squares). No secondary signals were observed when conditioning on rs11377254.

**Supplementary Figure 13**

Region plot for the *REL* locus. The credible set for RA contains a single variant with strong posterior (rs35149334), but shows no association in T1D. Conditioning on rs35149334, we do not observe a secondary association.

**Supplementary Figure 14**

Region plot for the *IL2RA* locus. The credible set for T1D contains two variants, with rs61839660 having the largest posterior (0.85). When performing conditional analysis, a secondary association is observed from rs4747846, a tertiary association from rs41295159, and finally, a quaternary association from rs704778. B) Pairwise exhaustive analysis in T1D shows that there are 0 pairs with a lower association p-value than rs61839660 + rs474846. C) Haplotype analysis suggests independent and opposite effects from haplotypes carrying rs681839660 and rs706778 alternate alleles.

**Supplementary Figure 15**

A) Region plots for the *CD28/CTLA4* locus: rs3087243, near <u>*CTLA4,*</u> has an increased posterior in the combined analysis compared with T1D, indicating a shared effect. In RA, rs117701653, near *CD28*, has the highest posterior. Both rs3087243 and rs11701653 are independently associated with RA, but not T1D. B) Exhaustive pairwise analysis for RA shows that the rs117701 653+rs3087243 pair has the strongest association for RA, but not T1D. C) Left to right: specific band in EMSA for rs117701653 C allele can be competed away using non-labeled A probe, indicating specific binding for A allele as well. Dose titration of labeled C and A allele probes (quantities in fmol) indicates that A allele also shows allele specific binding at higher probe quantities. EMSA in THP-1 monocyte cells does not show band for specific binding that is visible in Jurkat T cells for the rs117701653 C allele. EMSA for rs55686954 shows allele specific binding for the A allele. D) When performing a luciferase assay on rs117701653 and rs55686954, we observe allele specific enhancer activity for rs117701653 but not rs55686954.

**Supplementary Figure 16**

Promoter capture hi-C plots for the *CD28/CTLA4*, *TNAIP3* and *MEG3* loci show multiple contacts between bait sequences containing potential causal variants and downstream genomic regions. Figures adapted from http://www.chicp.org/

**Supplementary Figure 17**

Region plot for the *TNFAIP3* locus showing (from top to bottom) genes, posterior probabilities, and association p-values. The credible set for RA contains 8 variants, including indel rs35926684 (indicated by squares). No significant association was observed for T1D. When conditioning on rs35926684, a suggestive secondary signal was observed from rs58721818. B) Exhaustive pairwise testing shows that there are 6 pairs having a stronger association with RA than rs35926684 + rs58721818, with rs6920220 + rs58721818 showing the strongest association. C) Left to right: EMSA dose titration of labeled G and GA allele probes for rs355926684 (quantities in fmol) indicates that G allele also shows allele specific binding at higher probe quantities. Specific binding for the GA allele is not observed in THP-1 monocyte cells. EMSA in Jurkat cells for rs6920220 does not indicate specific binding.

**Supplementary Figure 18**

A) Region plot for the *MEG3* locus showing (from top to bottom) genes, posterior probabilities, and association p-values. We observe two variants in the credible set (rs56994090 and indel rs34552516; indicated by squares) for T1D, but no association in RA. We did not observe secondary signals when conditioning on rs56994090. B) EMSA in Jurkat T cells and THP-1 monocyte cells, shows no specific binding for the TC allele of rs34552516.

## Supplementary Tables

**Supplementary Table 1**

Overview of the cases and controls for each of the cohorts included in this study.

**Supplementary Table 2**

List of ImmunoChip regions, and regions with significant associations with RA or T1D published in previous studies.

**Supplementary Table 3**

Statistics for variants called from targeted sequencing experiment (MAF > 1%).

**Supplementary Table 4**

Imputation accuracy as determined by correlating imputed genotype dosages with genotypes called from targeted sequencing experiment for variants that are both present and absent on ImmunoChip.

**Supplementary Table 5**

Differences between imputation reference panels, by testing the difference in imputation accuracy (t-test).

**Supplementary Table 6**

Number of variants used as input for imputation, and output of imputation, at different imputation quality (INFO) score and allele frequency thresholds for each imputation reference panel.

**Supplementary Table 7**

Comparison of results presented in Okada et al. with the RA association analysis, for regions significant in this study. For each study, we compared the strongest association per region. Gt: genotyped variant.

**Supplementary Table 8**

Comparison of results presented in Onengut-Gumuscu et al. with the T1D association analysis, for regions significant in this study. For each study, we compared the strongest association per region. Gt: genotyped variant.

**Supplementary Table 9**

Correlations between RA and T1D association statistic Z-scores for the 76 tested loci.

**Supplementary Table 10**

Comparison of results from the combined analysis with the RA association analysis, for regions significant in this study. For each analysis, we compared the association with the strongest association per region. Gt: genotyped variant.

**Supplementary Table 11**

Comparison of results from the combined analysis with the T1D association analysis, for regions significant in this study. For each analysis, we compared the association with the strongest association per region. Gt: genotyped variant.

**Supplementary Table 12**

90% credible sets identified in this study: association results for regions that have ≤ 10 variants in the 90% credible set and are significant in either RA, T1D, or the combined analysis.

**Supplementary Table 13**

Conditional analysis results for the RA, T1D and combined analysis.

**Supplementary Table 14**

Haploreg annotations for candidate variants.

**Supplementary Table 15**

Overlap DNAse-I, H3K4me3, ChromHMM enhancers and ChromHMM promoters for both immune cell type groups and other cell types.

**Supplementary Table 16**

Overlap of credible sets with ATAC-seq peaks called from time course experiment in CD4+ T cells.

**Supplementary Table 17**

eQTL overlap for the 90% credible sets in the RA, T1D, and combined analysis.

**Supplementary Table 18**

Oligonucleotide probes used during EMSA analysis.

**Supplementary Table 19**

Primers used for cloning Luciferase assay plasmids.

## References

1. Maller, J. B. et al. Bayesian refinement of association signals for 14 loci in 3 common diseases. Nat. Genet. 44, 1294–301 (2012).

2. Wakefield, J. A Bayesian Measure of the Probability of False Discovery in Molecular Genetic Epidemiology Studies (DOI:10.1086/519024). Am. J. Hum. Genet. 83, 424 (2008).

3. Eyre, S. et al. High-density genetic mapping identifies new susceptibility loci for rheumatoid arthritis. Nat. Genet. 44, 1336–40 (2012).

4. Onengut-Gumuscu, S. et al. Fine mapping of type 1 diabetes susceptibility loci and evidence for colocalization of causal variants with lymphoid gene enhancers. Nat. Genet. 47, 381–386 (2015).

5. Klareskog, L., Catrina, A. I. & Paget, S. Rheumatoid arthritis. Lancet 373, 659–672 (2009).

6. Palmer, J. P. et al. Insulin antibodies in insulin-dependent diabetics before insulin treatment. Science 222, 1337–9 (1983).

7. Baekkeskov, S. et al. Identificationof the 64K autoantigen in insulin-dependent diabetes as the GABA-synthesizing enzyme glutamic acid decarboxylase. Nature 347, 151–156 (1990).

8. Okada, Y. et al. Genetics of rheumatoid arthritis contributes to biology and drug discovery. Nature 506, 376–81 (2014).

9. Huang, H. et al. Association mapping of inflammatory bowel disease loci to single variant resolution. bioRxiv 28688 (2015). doi:10.1101/028688

10. Gaulton, K. J. et al. Genetic fine mapping and genomic annotation defines causal mechanisms at type 2 diabetes susceptibility loci. Nat. Genet. 47, 1415–1425 (2015).

11. Farh, K. K. et al. Genetic and epigenetic fine mapping of causal autoimmune disease variants. Nature 518, 337–343 (2015).

12. Trynka, G. et al. Chromatin marks identify critical cell types for fine mapping complex trait variants. Nat. Genet. 45, 124–30 (2013).

13. McCarthy, S. et al. A reference panel of 64,976 haplotypes for genotype imputation. Nat. Genet. 48, 1279–1283 (2016).

14. Abecasis, G. R. et al. A map of human genome variation from population-scale sequencing. Nature 467, 1061–73 (2010).

15. Begovich, A. B. et al. A missense single-nucleotide polymorphism in a gene encoding a protein tyrosine phosphatase (PTPN22) is associated with rheumatoid arthritis. Am. J. Hum. Genet. 75, 330–7 (2004).

16. Bottini, N. et al. A functional variant of lymphoid tyrosine phosphatase is associated with type I diabetes. Nat. Genet. 36, 337–338 (2004).

17. Diogo, D. et al. TYK2 protein-coding variants protect against rheumatoid arthritis and autoimmunity, with no evidence of major pleiotropic effects on non-autoimmune complex traits. PLoS One 10, e0122271 (2015).

18. Zochling, J. et al. An Immunochip-based interrogation of scleroderma susceptibility variants identifies a novel association at DNASE1L3. Arthritis Res. Ther. 16, (2014).

19. Al-Mayouf, S. M. et al. Loss-of-function variant in DNASE1L3 causes a familial form of systemic lupus erythematosus. Nat. Genet. 43, 1186–1188 (2011).

20. Ueki, M. et al. Caucasian-specific allele in non-synonymous single nucleotide polymorphisms of the gene encoding deoxyribonuclease I-like 3, potentially relevant to autoimmunity, produces an inactive enzyme. Clin. Chim. Acta 407, 20–24 (2009).

21. Fortune, M. D. et al. Statistical colocalization of genetic risk variants for related autoimmune diseases in the context of common controls. Nat. Genet. 47, 839–46 (2015).

22. Aguet, F. et al. Local genetic effects on gene expression across 44 human tissues. bioRxiv (2016).

23. Javierre, B. M. et al. Lineage-Specific Genome Architecture Links Enhancers and Non-coding Disease Variants to Target Gene Promoters. Cell 167, 1369–1384.e19 (2016).

24. Tsoi, L. C. et al. Identification of 15 new psoriasis susceptibility loci highlights the role of innate immunity. Nat. Genet. 44, 1341–1348 (2012).

25. Jostins, L. et al. Host–microbe interactions have shaped the genetic architecture of inflammatory bowel disease. Nature 491, 119–124 (2012).

26. Beecham, A. H. et al. Analysis of immune-related loci identifies 48 new susceptibility variants for multiple sclerosis. Nat. Genet. 45, 1353–1360 (2013).

27. Lessard, C. J. et al. Variants at multiple loci implicated in both innate and adaptive immune responses are associated with Sjögren’s syndrome. Nat. Genet. 45, 1284–1292 (2013).

28. Cordell, H. J. et al. International genome-wide meta-analysis identifies new primary biliary cirrhosis risk loci and targetable pathogenic pathways. Nat. Commun. 6, 8019 (2015).

29. Bentham, J. et al. Genetic association analyses implicate aberrant regulation of innate and adaptive immunity genes in the pathogenesis of systemic lupus erythematosus. Nat. Genet. 47, 1457–1464 (2015).

30. Trynka, G. et al. Dense genotyping identifies and localizes multiple common and rare variant association signals in celiac disease. Nat. Genet. 43, 1193–201 (2011).

31. McGovern, A. et al. Capture Hi-C identifies a novel causal gene, IL20RA, in the pan-autoimmune genetic susceptibility region 6q23. Genome Biol. 17, 212 (2016).

32. Zhou, Y. et al. Activation of p53 by MEG3 Non-coding RNA. J. Biol. Chem. 282, 24731–24742 (2007).

33. Wallace, C. et al. The imprinted DLK1-MEG3 gene region on chromosome 14q32.2 alters susceptibility to type 1 diabetes. Nat. Genet. 42, 68–71 (2010).

34. Purcell, S. et al. PLINK: a tool set for whole-genome association and population-based linkage analyses. Am. J. Hum. Genet. 81, 559–75 (2007).

35. Browning, B. L. & Browning, S. R. Genotype Imputation with Millions of Reference Samples. Am. J. Hum. Genet. 98, 116–126 (2016).

36. Loh, P.-R. et al. Reference-based phasing using the Haplotype Reference Consortium panel. Nat. Genet. 48, 1443–1448 (2016).

37. Delaneau, O., Marchini, J. & Zagury, J.-F. A linear complexity phasing method for thousands of genomes. Nat. Methods 9, 179–181 (2011).

38. Durbin, R. Efficient haplotype matching and storage using the positional Burrows-Wheeler transform (PBWT). Bioinformatics 30, 1266–72 (2014).

39. Fuchsberger, C., Abecasis, G. R. & Hinds, D. A. minimac2: faster genotype imputation. Bioinformatics 31, 782–4 (2015).

40. Li, H. & Durbin, R. Fast and accurate short read alignment with Burrows-Wheeler transform. Bioinformatics 25, 1754–60 (2009).

41. Van der Auwera, G. A. et al. From FastQ data to high confidence variant calls: the Genome Analysis Toolkit best practices pipeline. Curr. Protoc. Bioinformatics 43, 11.10.1-33 (2013).

42. Powell, J. E., Visscher, P. M. & Goddard, M. E. Reconciling the analysis of IBD and IBS in complex trait studies. Nat. Rev. Genet. 11, 800–805 (2010).

43. Zhernakova, D. V et al. Identification of context-dependent expression quantitative trait loci in whole blood. Nat. Genet. (2016). doi:10.1038/ng.3737

44. Raj, T. et al. Polarization of the Effects of Autoimmune and Neurodegenerative Risk Alleles in Leukocytes. Science (80-.). 344, 519–523 (2014).

45. Ernst, J. & Kellis, M. ChromHMM: automating chromatin-state discovery and characterization. Nat. Methods 9, 215–216 (2012).

46. Roadmap Epigenomics Consortium, A. et al. Integrative analysis of 111 reference human epigenomes. Nature 518, 317–30 (2015).

47. Ward, L. D. & Kellis, M. HaploReg v4: systematic mining of putative causal variants, cell types, regulators and target genes for human complex traits and disease. Nucleic Acids Res. 44, D877–D881 (2016).

48. Buenrostro, J. D., Giresi, P. G., Zaba, L. C., Chang, H. Y. & Greenleaf, W. J. Transposition of native chromatin for fast and sensitive epigenomic profiling of open chromatin, DNA-binding proteins and nucleosome position. Nat. Methods 10, 1213–8 (2013).

49. Zhang, Y. et al. Model-based analysis of ChIP-Seq (MACS). Genome Biol. 9, R137 (2008).

